# IKK2/β mediated phosphorylation of transcription factor Ets2 at site(s) distal to DNA binding domain negatively modulates its DNA binding activity

**DOI:** 10.64898/2026.08.31.747788

**Authors:** Deeparna Sutradhar, Anna Rose Antony, Afreen Haque, Prateeka Borar, T Pallavi Rao, Swasti Raychaudhuri, Prasun Kumar, Smarajit Polley

**Affiliations:** Department of Biological Sciences, Bose Institute, Unified Academic Campus, EN 80, Sector V, Bidhan Nagar, Kolkata 700091, West Bengal, India; Department of Biological Sciences and Engineering, Indian Institute of Technology Palakkad, Kanjikode, Palakkad 678623, Kerala, India; CSIR-CCMB, Hyderabad, Telengana, India; EMBL, Heidelberg

## Abstract

Transcription factor Ets2 coalesce with the NF-κB pathway to regulate gene expression in specific signaling contexts. IKK2/β-mediated phosphorylation events critically regulate the NF-κB pathway. However, any link between Ets2 and IKK2 remains elusive. Here we report Ets2 as a direct substrate of IKK2. *In-vitro* kinase assays using deletion constructs, high resolution MS-MS and site directed mutagenesis identified S295 as a prominent phosphorylation site distal to the DNA binding domain, substitution of which to phosphor-mimetic Glutamate triggers further phosphorylation of Ets2. MD simulations clearly indicate conformational constriction of the otherwise disordered N-terminal region and inhibition of DNA binding activity upon phosphorylation, which was further confirmed by Electrophoretic mobility shift assays. Our results uncover a phosphoregulatory connection between Ets2 and IKK2.

## Introduction

Phosphorylation is known to be the most common post-translational modification (PTM) in proteins. Phosphorylation mediated regulation of transcription factor activity through modulation of its DNA interaction is well studied [1–7]. IKK2 mediated phosphorylation of canonical substrate IκBα leads to NF-κB activation, thus making IKK2 a pivotal player controlling inflammatory responses [8,9]. IKK2 has also been shown to phosphorylate and regulate activities of several other proteins such as β-catenin [10], SRC-3 [11], TSC1 [12], FOXO3a [13], p65 [14,15] and p53 [16]. Inflammatory pathways also involve multiple transcription factors that work in coordination [17]. However, whether IKK2 targets other transcription factors involved in inflammation remains less explored.

E26 transformation-specific 2 (Ets2), one of the founder members of the ETS family of transcription factors is known to participate in inflammation regulating pro-inflammatory cytokines like TNFα, IL1β and IL6[18–23]. Recent studies have shown Ets2 to be a central regulator of inflammatory bowel disease (IBD) [24–27]. Ets2 supresses MAPK/NF-κB pathway in LPS or VSV stimulated inflammation [18]. It is a core regulator of inflammatory macrophage polarisation through TLR4/ NF-κB signalling pathways in ulcerative colitis [22] [28]. It is also known to aggravate allergic airway inflammation in asthma[29]. Being linked to chronic inflammation, Ets2 can also contribute to inflammation associated tumorigenesis [30–34].

Transcription factors are amenable to regulatory post translational modifications[35–37]. Information curated from several papers report a total of 35 PTMs in PhosphoSitePlus database for Ets2 [38]. Although majority of the reported PTMs are phosphorylation, the kinases responsible for phosphorylating these residues upstream remains unidentified. The functional outcome of such phosphorylation events on DNA binding properties of Ets2 hence remains under explored. On the hand, phosphorylation mediated inhibition of DNA-binding by Ets1 is well established in the literature.

IKKs are critical regulators of inflammation and related signalling events, and Ets2 being a key transcription factor in inflammatory pathways, we explored the possibility of any cross-regulation between them. In this study we have investigated whether IKK2 can phosphorylate Ets2 and identify the residues involved. Using a combination of biochemical approaches including in-vitro kinase assays, DNA-binding assays, structural modelling and all-atom molecular dynamics (MD) simulations, we determine the significance of the key residues involved in the modification, examine how phosphorylation alters the conformational dynamics of the N-terminal region (NTR) of Ets2, and elucidate how these changes impact its DNA binding. These findings help uncover the structural and biochemical mechanism of phosphorylation-driven regulation of transcription factor Ets2.

## Results

### IKK2 phosphorylates Ets2 *in-vitro*

We used a pre-defined phosphorylation site motif of IKK2 for screening potential substrate recognition sequence in Ets2 (Figure 1A)[16]. A sequence-based search within Ets2 identified a region containing candidate serine residues that aligns well with the consensus sequence. The identified region exhibits high sequence similarity with the motif present in IκBα, a well-known substrate of IKK2 (Figure 1B). The sequence alignment using Weblogo [39] in Figure 1C shows this phosphorylation motif in Ets2 being evolutionary conserved across species. These results indicate that Ets2 might be a substrate of IKK2.

**Figure 1:**
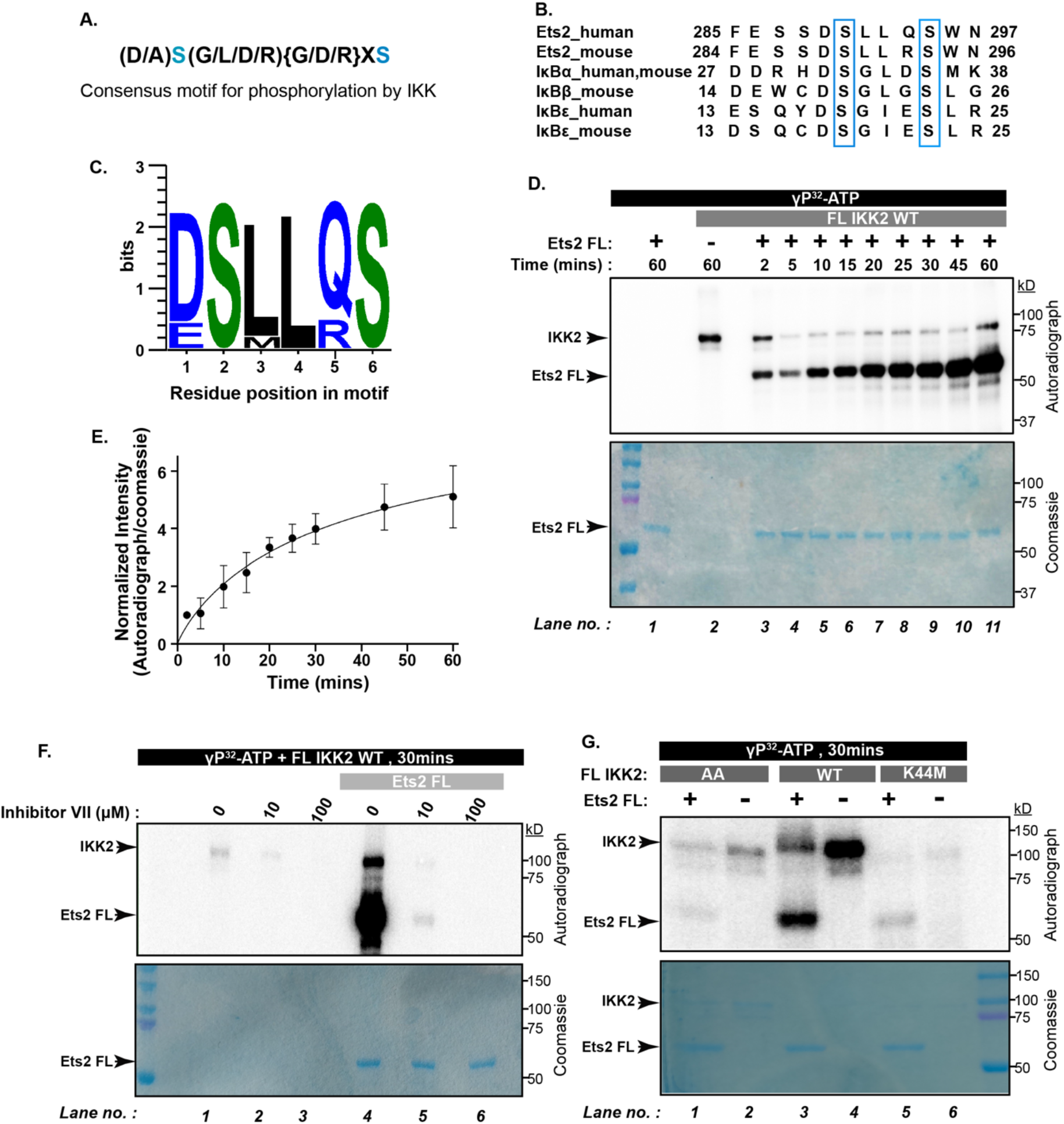
IKK2 phosphorylates Ets2 in-vitro. **(A)** The IKK2 consensus phosphorylation motif showing the acceptable amino acids in squared brackets () in position 1 and 3 and the unacceptable amino acids in curly brackets {} in position 4. Phosphorylatable Serine residues are shown in blue. **(B)** Alignment of peptide sequences containing IKK2 phosphorylation sites. Serine residues in canonical substrates that are phosphorylated by IKK complex are shown in blue boxes. **(C)** Sequence logo plot using WebLogo showing the conservation of IKK2 consensus phosphorylation motif in Ets2 across species. **(D)** *In-vitro* radioactive kinase assay showing the phosphorylation of Ets2 by IKK2 at different time points (n=3). **(E)** The normalized intensity (Autoradiograph/Coomassie) was plotted against time. The data values are shown as mean ± s.d. of normalized intensity values from three independent measurements (n=3). **(F)** Effect of Inhibitor VII specific for IKK2 on kinase autophosphorylation and Ets2 substrate phosphorylation at different inhibitor concentrations in a radioactive *in vitro* kinase assay. This assay was performed twice. **(G)** Comparison of substrate phosphorylation and autophosphorylation activities of wild type and different catalytic mutants of IKK2 in an *in vitro* radioactive kinase assay performed with or without FL Ets2 as the substrate. This assay was performed twice.

**Figure S1:**
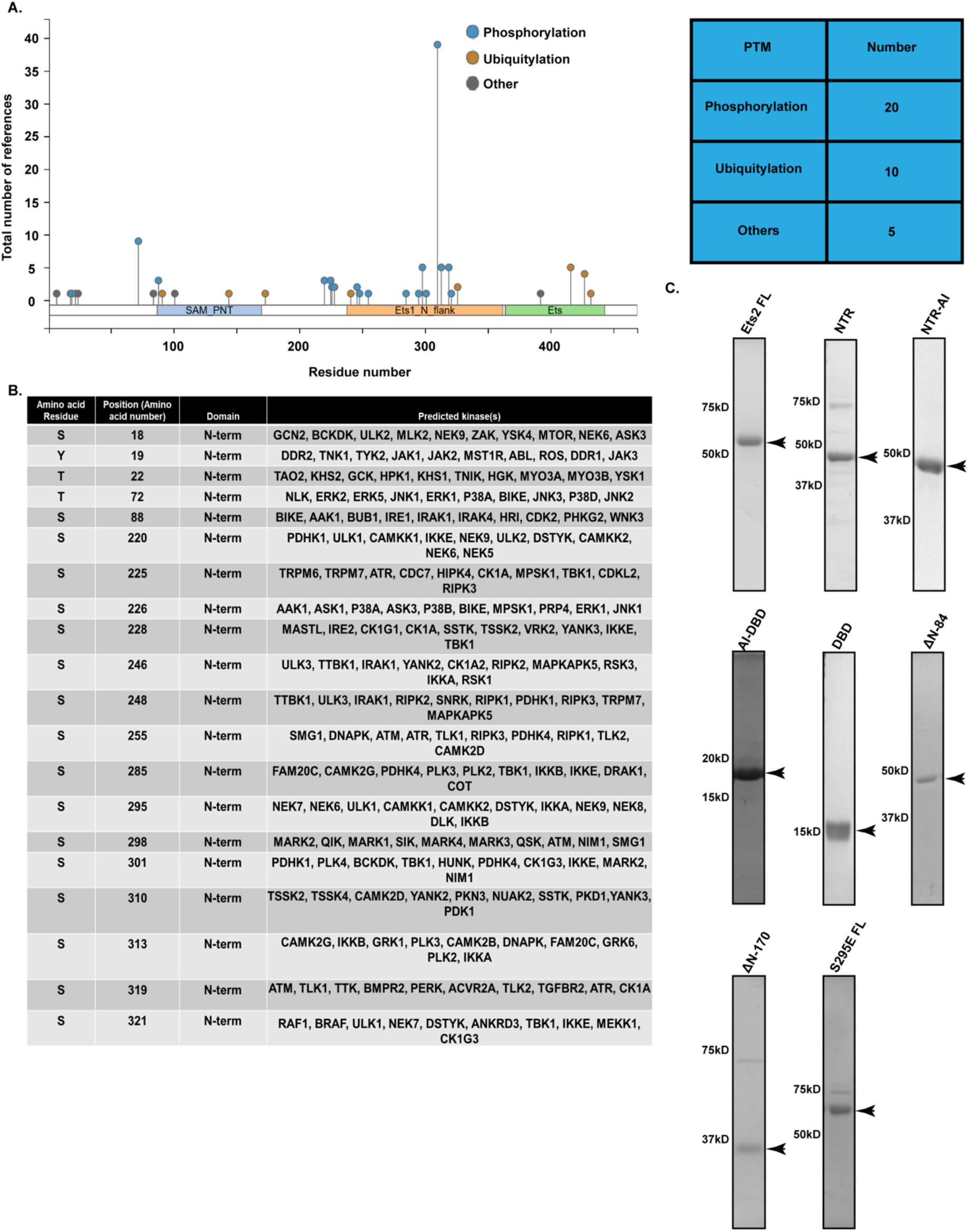
**(A)** Reported PTMs of Ets2 in PhosphositePlus website. **(B)** Predicted kinases for the phosphorylation of STY residues of Ets2 using web tools. **(C)** Coomassie stained SDS-PAGE showing all purified proteins used in this study.

Out of the many phosphorylation sites reported in PhosphoSitePlus [38] database for Ets2 (Figure S1A), only three kinases – CAMK2A, CDK10 and HRas are known to phosphorylate 6 serine residues[40–44]. For the remaining sites, the responsible kinases remain unidentified and not reported in any publicly available database[45]. We next performed a prediction of putative kinases for the phosphorylated residues using The Kinase Library and PhosphoNET (Figure S1B). The IKK kinases have been predicted for some residues prompting us to investigate further.

The ability of IKK2 to phosphorylate recombinant full-length human Ets2 *in-vitro* was tested using γ-P^32^-ATP as the phosphate donor. Figure 1D shows that IKK2 could robustly phosphorylate Ets2 and the phosphorylation signal showed a time-dependent increase (Figure 1E).

Next, we used IKK-specific ATP-competitive inhibitor, Calbiochem Inhibitor VII to confirm such phosphorylation was indeed IKK2-mediated using similar *in-vitro* kinase assay, wherein both IKK2-autophosphorylation (Figure 1F, *lane 1-3*) and Ets2 phosphorylation (Figure 1F, *lane 4-6*) were diminished as a function of inhibitor concentration. To further confirm these observations, we used two mutants of IKK2, K44M and S177A/S181A. In the IKK2 K44M mutant highly conserved ATP-anchoring K44 is mutated to M that is deleterious to kinase activity, whereas in IKK2 AA two activation loop Ser residues, phosphorylation of which are known hallmarks of active IKK2, are mutated to phosphoablative Ala. Both these mutants of IKK2 (Figure 1G, lanes 1, 2 and 5, 6) showed severely compromised autophosphorylation and Ets2 phosphorylation in *in vitro* kinase assay as compared to the wild type (WT) IKK2 (Figure 1G, lanes 3, 4). The above sets of experiments confirm that Ets2 is a substrate of IKK2 that is phosphorylated by the canonical active site of the kinase and activation loop phosphorylation of IKK2 is required to phosphorylate Ets2.

### IKK2 phosphorylates Ets2 predominantly at its N-terminal Serine Rich Region but not at autoinhibition domain or DNA binding domain

To map the residue(s) phosphorylated by IKK2, we created seven deletion constructs of Ets2 and purified those truncated proteins for *in vitro* kinase assays (Figure 2A). We observed that the NTR of Ets2 was more efficiently phosphorylated by IKK2 than the DNA binding domain (DBD) or the DBD connected to the autoinhibitory domain (AI-DBD). Interestingly, the presence of NTR (AI-NTR) or any part of NTR (ΔN-84 and ΔN-170) showed efficient phosphorylation of those constructs. This experiment strongly suggests that Ets2 gets phosphorylated by IKK2 at multiple residues (Figure 2B).

**Figure 2:**
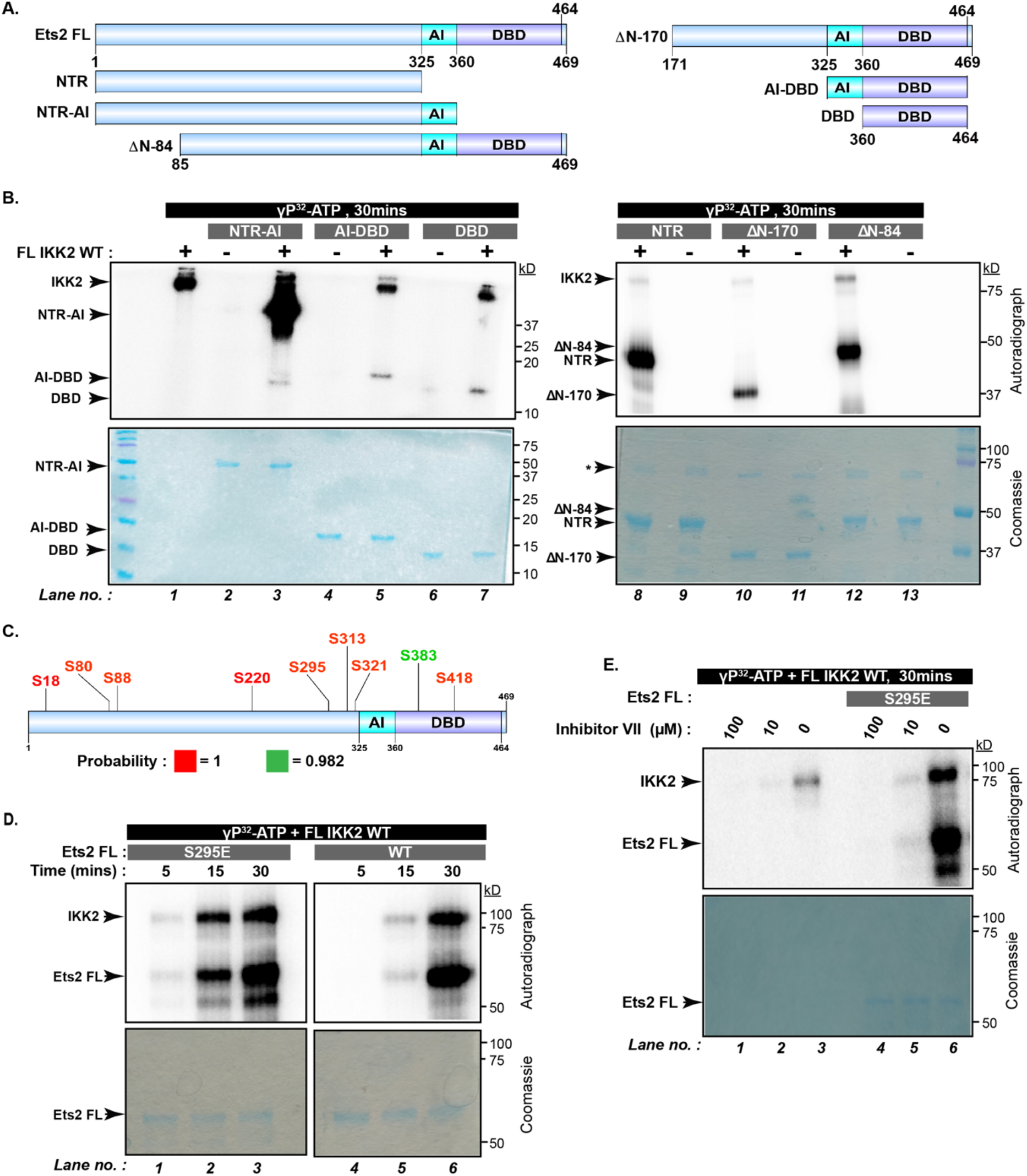
IKK2 phosphorylates Ets2 primarily at the NTR. **(A)** Schematic representation of the domain organisation of Ets2 FL depicting autoinhibition domain (AI) and DNA binding ETS domain (DBD) based on X-ray crystal structure (pdb id 4bqa) and N-terminal region (NTR). All truncated versions of Ets2 used in this study are also depicted schematically. **(B)** Radioactive kinase assay conducted to check phosphorylation of Ets2 deletion constructs by FL IKK2 WT. This assay was performed twice. **(C)** Phosphorylation sites identified by LC-MS/MS experiments are shown on cartoon representation of Ets2. Residues coloured in Red have been identified with a probability = 1 and residue coloured in green has a probability of 0.982. **(D)** Comparison of substrate phosphorylation between FL Ets2 WT and S295E with FL IKK2 WT at different time points using *in-vitro* radioactive kinase assay. **(E)** Effect of Inhibitor VII on FL IKK2 WT autophosphorylation and Ets2 S295E substrate phosphorylation at different inhibitor concentrations in a radioactive *in vitro* kinase assay. This assay was performed twice.

**Figure S2:**
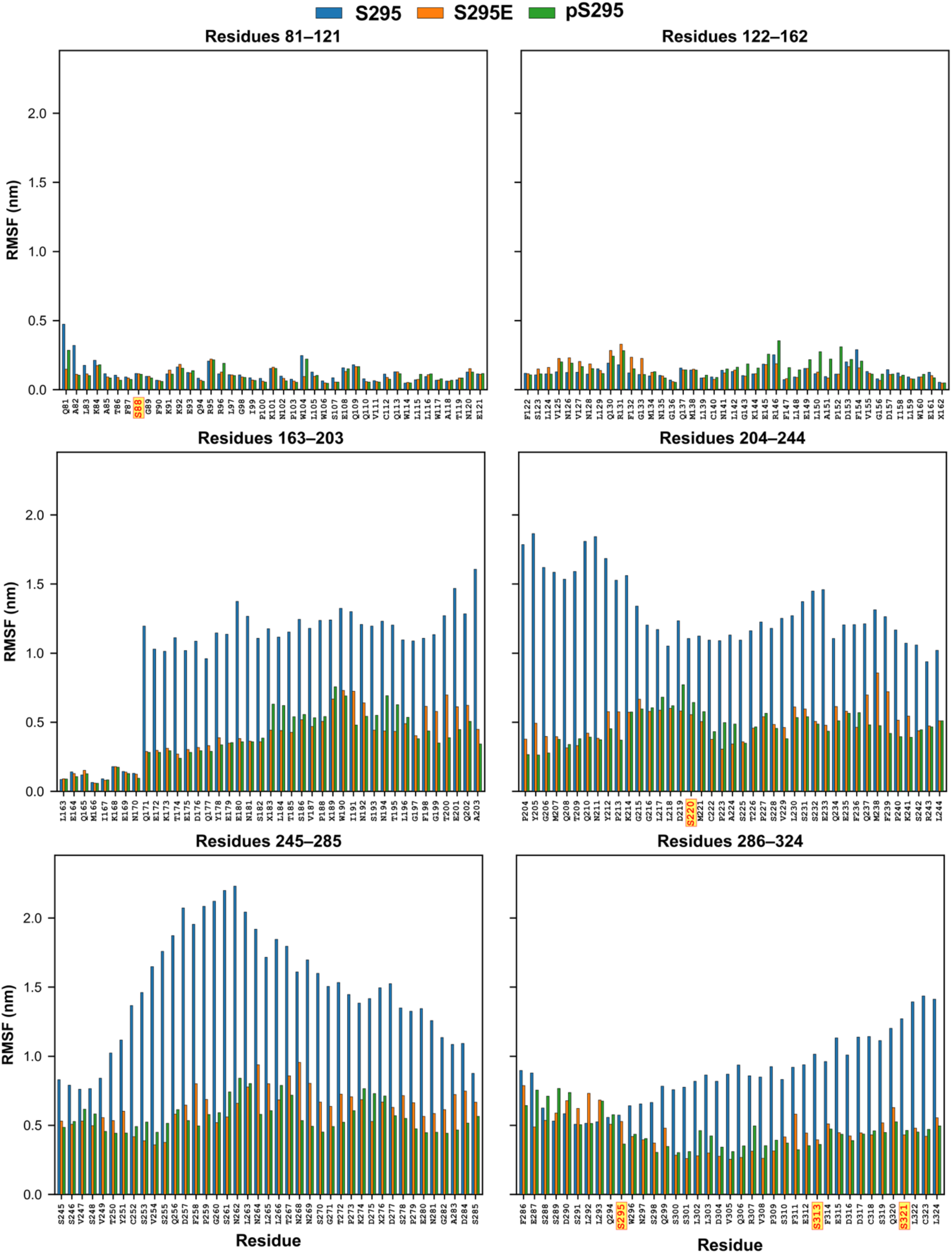
Residue-wise RMSF comparison of N-terminal residues to residue number 295. RMSF profiles of the residues that are N-terminal to residue 295 in the WT S295 (blue), S295E (orange), and pS295 (green). Phosphorylatable serine residues are highlighted in yellow.

To identify residues phosphorylated by IKK2, we performed LC-MS/MS experiment on an Orbitrap. Mass spectrometry results revealed that Ets2 undergoes phosphorylation at multiple sites by IKK2. We only considered residues with probability of phosphorylation greater than 0.97 (Figure 2C). Seven phosphorylated serine residues: S18, S80, S88, S220, S295, S313, S321 were located within the NTR explaining the extensive phosphorylation that was seen in the *in-vitro* kinase assay in Figure 2B and 2C. Two other phosphorylated residue S383 and S418 were situated in the DBD of Ets2 giving possible explanation for the less substrate (AI-DBD or DBD) phosphorylation seen in Figure 2B.

Based on the sequence comparison of IKK2-recognition motif on Ets2 with the same on IκBα (Figure 1A), we generated phosphor-mimetic (S295E) mutant of Ets2. Interestingly, mutation of S295 to phosphomimmetic E resulted in enhanced phosphorylation compared to Ets2 WT (Figure 2D). We surmise that S295 phosphorylation is critical to multisite phosphorylation of Ets2 that triggers phosphorylation of other site(s). Abrogation of phosphorylation of Ets2 S295E in presence of IKK2-specific inhibitor, Inhibitor VII, strongly suggests that multisite phosphorylation on S295E is also mediated by IKK2 and not any other spurious kinase activity(Figure 2E).

### Phosphorylated S295 anchors the NTR to the DNA-binding face of the DBD

To characterize the spatial behavior of residue 295 in different phospho-states, we performed 500 ns all-atom MD simulations of three states, namely (a) unmodified S295; (b) phosphomimmetic S295E and phosphoserine pS295, on a construct lacking the first 80-amino acids (ΔN-80). In the RMSF analyses of upstream residues of S295 (Figure S2), the conformational flexibility is significantly reduced in both S295E and pS295 compared to S295, indicating that S295 modification puts an intramolecular restraint on the disordered region. We also explored its occupancy by analyzing the distribution of positions sampled during the trajectory (Figure 3A and 3B). In the Ets2 WT simulation, residue 295 is found in a single, relatively confined spatial envelope, consistent with a mobile but loosely tethered residue in the NTR (Figure 3A and column 1). In the S295E simulation, the occupancy cloud is more constrained, indicating more restricted positional sampling (Figure 3A and column 2). In contrast, the pS295 simulation shows a bimodal distribution of occupancies with two spatial clusters, suggesting that the highly electronegative phosphoserine at position 295 interconverts between two preferred conformational sub-states (Figure 3A and column 3). Such bimodal behavior is lacking for the simpler phosphomimmetic glutamic acid, showing that the multiply charged state of a real phosphoserine imparts unique conformational properties that the singly charged glutamic acid cannot replicate.

**Figure 3:**
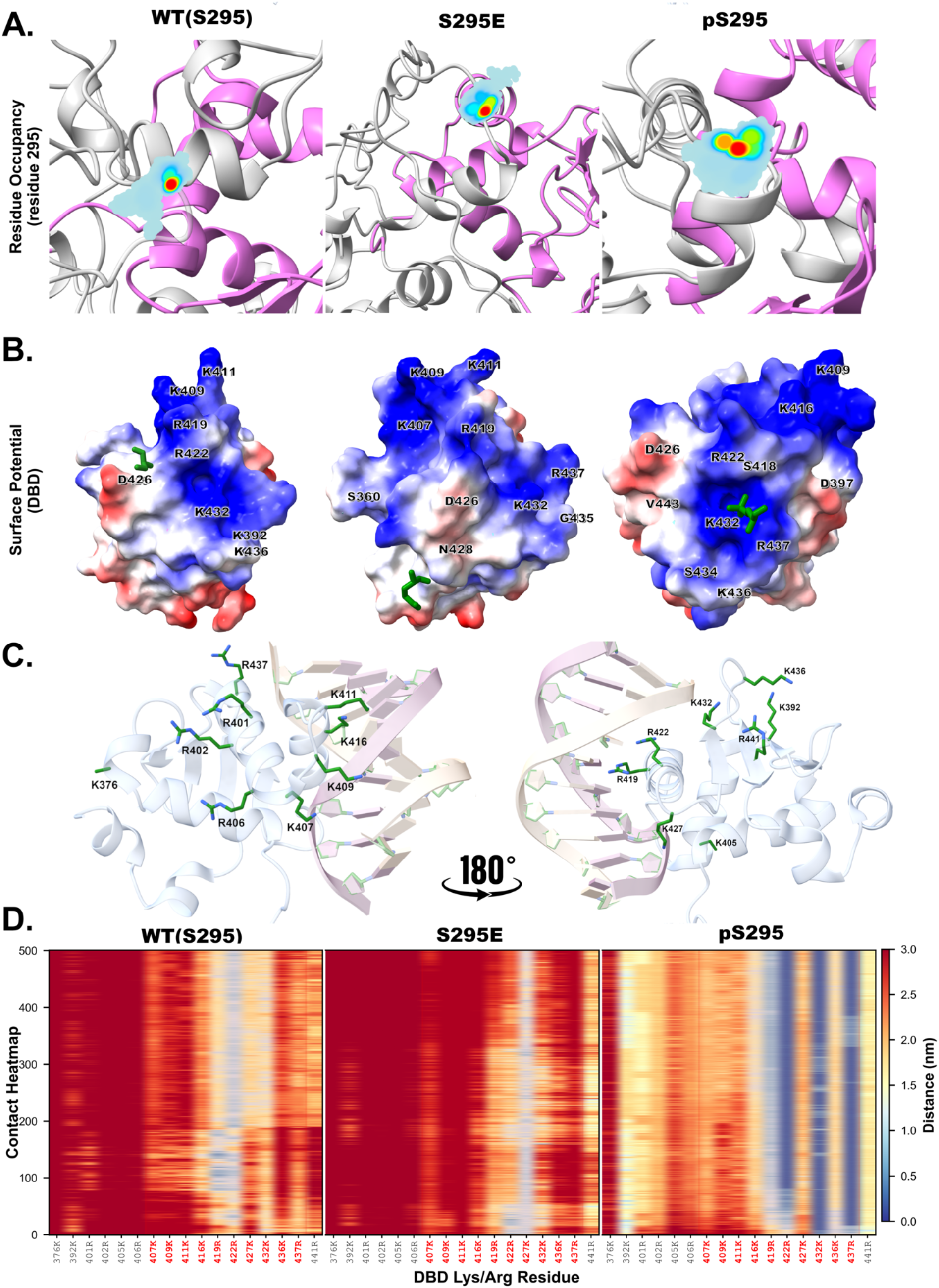
MD data analyses of residue 295 positioning and its interactions with the Ets2 DBD. **(A)** Residue 295 occupancy maps for the WT (S295), S295E, and pS295 states in the ΔN-80 construct. The DBD is shown in magenta and the remaining protein regions in gray. Cyan density surfaces represent the spatial distribution of residue 295, with warmer colors (yellow-red) indicating higher occupancy.(B) Electrostatic surface representation of the Ets2 DBD, colored from negative (red) to positive (blue) electrostatic potential. Residue 295 is shown in green sticks. A few residues of DBD are also labeled. **(C)** Positively charged Lys/Arg residues of the DBD mapped on the DNA-DBD complex (PDB ID: 4bqa). **(D)** Pairwise distance analysis between residue 295 and basic residues throughout the 500 ns MD simulations. DNA-binding basic residues are shown in red-colored text.

Though S295 is not a part of DBD and lies within the intrinsically disordered NTR, yet different phospho-states localise around the face of the DBD that interacts with DNA. This observation prompted us to investigate direct contacts with the positively charged lysine (K) and arginine (R) residues that constitute the DNA-binding surface of the DBD. We calculated the pairwise distances between residue 295 and all DBD Lys/Arg residues (K376, K392, R401, R402, K405, R406, K407, K409, K411, K416, R419, R422, K427, K432, K436, R437, R441)

(Figure 3C) along the entire 500 ns trajectory (Figure 3D). The resulting distance-maps reveal that the wild-type S295 is typically at longer distances from the majority of basic residues in the DBD, establishing only transient close contacts. In contrast, the pS295 simulation displays persistent close proximity (< 1.0 nm) with DNA binding basic residues of DBD (K392, R402, K416, K427, K432, K436, R437 and R441) over long trajectory windows. The phosphomimmetic S295E shows intermediate behavior with less frequent and shorter close contacts than pS295. Thus, the addition of a negative charge at position 295, particularly the high charge density of a phosphoserine induces an electrostatic interaction with the basic residues of the DBD. This electrostatic tethering of NTR–DBD is consistent with the RMSF reduction noted above; the intrinsically disordered NTR is conformationally restrained as pS295 intermittently tethers to DBD basic residues.

In total, the reduction of RMSF (Figure S2), the localization of residue 295 to the DNA-binding face (Figure 3B-D) and the close proximity to the DNA-binding basic residues (Figure 3D) point to a model where IKK2-mediated phosphorylation of S295 anchors the IDR on the DNA-binding surface, directly occluding DNA-access that may, in turn, reduce binding affinity.

### IKK2 mediated phosphorylation of Ets2 modulates DNA binding

Based on our computational analyses, we hypothesized that phosphorylation of Ets2 is likely to directly interfere with its DNA binding activity. We investigated effects of IKK2 mediated phosphorylation on Ets2 DNA binding, if any, using EMSA. To this end, we phosphorylated Ets2 recombinantly expressed and purified from *E. coli* with pure IKK2 (PEts2, pan-phopshorylated) and compared its DNA-binding activity with that of the unphosphorylated protein. We observed marked reduction in DNA binding in phosphorylated Ets2 (K_d_ ~0.9µM) compared to its unphosphorylated counterpart (K_d_ ~0.45 µM) (Figure 4A). We also purified the S295E phosphomimmetic version of Ets2 and performed the same assay. EMSA results show inhibited or reduced DNA binding by S295E relative to wild-type Ets2 (Figure 4B). S295E binds to DNA with a K_d_ of ~0.9 µM with a cooperativity feature that was not apparent in Ets2 WT. Interestingly, pan-phosphorylated P-Ets2 and S295E versions of Ets2 display similar DNA-binding affinity strongly suggesting that phosphorylation mediated reduction of Ets2-DNA binding emanates primarily from tethering of phosphorylated S295 onto its DNA-binding interface.

**Figure 4:**
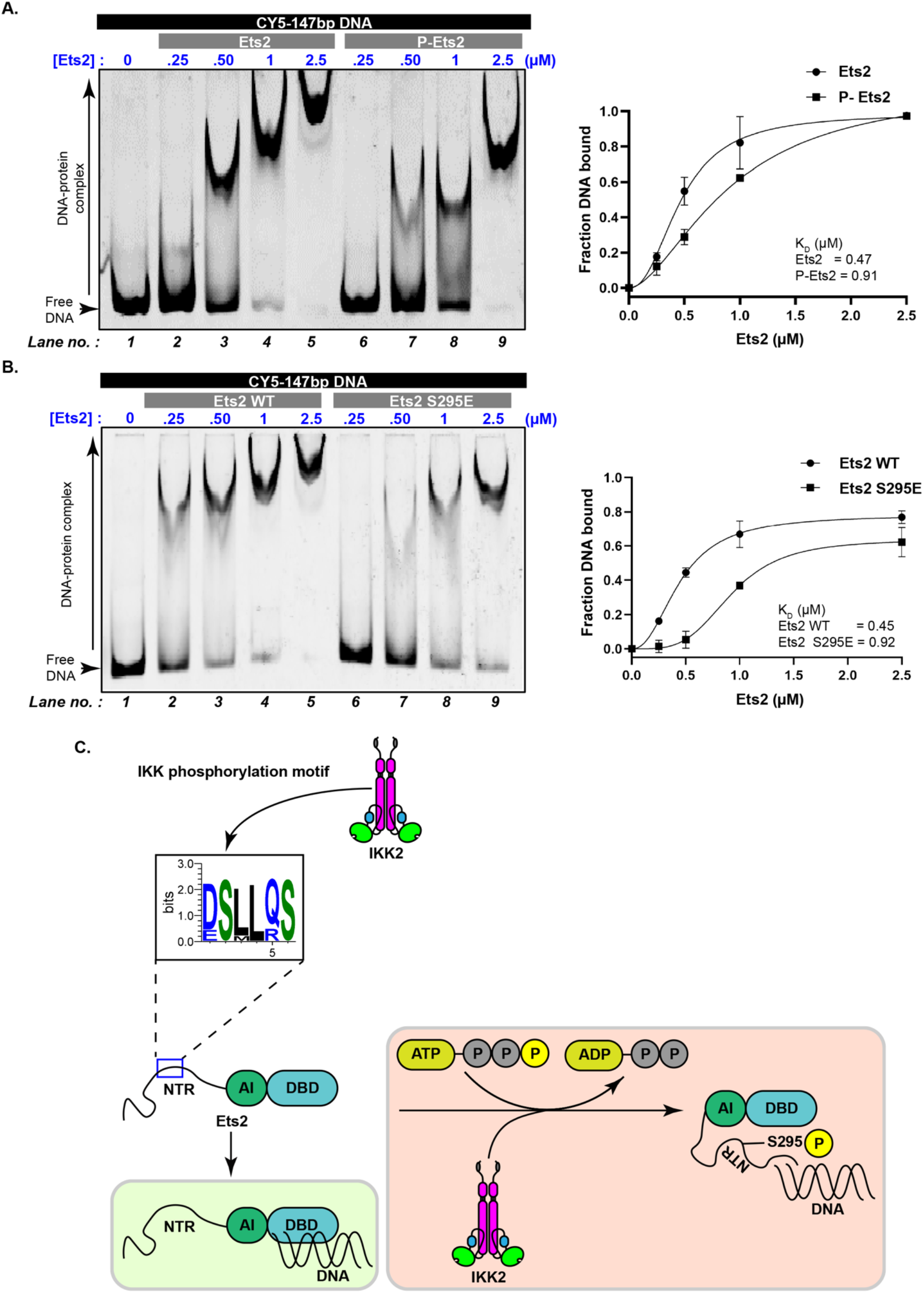
IKK2 mediated Ets2 phosphorylation inhibits DNA binding. **(A)** Electrophoretic mobility shift assay comparing Cy5-147bp DNA binding by Ets2 and P-Ets2 (left) and quantified (right) relative to the lane no. 1 in the EMSA gel. **(B)** EMSA comparing DNA binding by FL Ets2 WT and S295E (left) and quantified (right) relative to the lane no. 1 in the EMSA gel. The data values are shown as mean ± s.d. of normalized intensity values from three independent measurements (n=3). **(C)** Graphical representation illustrating IKK2 mediated phosphorylation of Ets2 at NTR residue S295 inhibits DNA binding.

## Discussion

Phosphorylation is one of the fundamental mechanisms to control activities of transcription factors [1]. Here, we report Ets2 transcription factor as a direct substrate of a Ser/Thr kinase IKK2. We found that IKK2 mediated phosphorylation of Ets2 negatively regulates its DNA-binding activity. Although primary target of IKK2 is the IκB-proteins to activate NF-κB pathway in response to a diverse array of stimuli [46], it can also directly phosphorylate transcription factor RelA, one of the five NF-κB subunits [14,15]. However, IKK2’s influence on other transcription factors remains somewhat under-appreciated, despite the fact that IKK2 is known to regulate transcription factors like p53 and FOXO3a. IKK2 mediated phosphorylation triggers Mdm2-independent degradation of p53 via β-TRCP1 [16], and subcellular localization of FOXO-3a [11,13]. Our findings therefore is a valuable addition to the known substrate landscape of IKK2.

We have identified a consensus IKK2-phosphorylation motif in Ets2 that is well in agreement with the phosphorylation motif found in canonical substrate of IKK2, IκBα. *In-vitro* kinase assays confirm Ets2 as a substrate of IKK2. Phosphorylation of Ets2 is mediated through the canonical active site of IKK2 as validated by the abolition of Ets2 phosphorylation by a specific IKK2 inhibitor and kinase-dead or activation loop mutants of IKK2. Taken together, these observations established Ets2 as a hitherto unknown direct substrate of IKK2.

Experiments to map site(s) of phosphorylation using truncated Ets2 proteins revealed that those sites are mainly located in the NTR. Of note, many ETS transcription factors harbor serine rich sequences which are preferred sites of phosphorylation by different kinases[47]. Such regions are often intrinsically disordered, making them more accessible and amenable to post-translational modifications[48–50]. Ets2 NTR also contains serine rich regions (SRR) explaining the preference of NTR over other domains to be modified by IKK2 [51]. We found that S295 within the NTR and distal to DBD as one of the primary candidate residues for phosphorylation[45]. S295E mutant exhibited enhanced phosphorylation compared to Ets2 WT protein. We speculate that phosphorylation of S295 may prime Ets2 for multisite phosphorylation in a coordinated manner, precise mechanism and role of which are yet to be deciphered.

MD simulations of different states of Ets2 with respect to S295 phosphorylation, namely unphosphorylated or S295, phosphorylated or pS295 and phosphomimmetic or S295E provide important structural insights of Ets2’s DNA-binding activity and its regulation through allosteric means. The NTR containing S295 shows high conformational entropy in the wild-type state, a feature of intrinsically disordered regions that makes S295 structurally available to upstream kinases like IKK2, in line with the serine-rich NTR composition discussed above. S295 in the phosphorylated states comes in contact with positively charged residues on the DNA-binding face of the DBD that causes considerable constriction of the conformational space of the otherwise disordered NTR region. Positional occupancy analysis clearly showed that phosphorylation or phosphomimmetic substitution of S295 significantly alters its positional distribution. These analyses further revealed the bimodal occupancy of pS295 that is in striking contrast to unimodal distribution in the unphosphorylated or phosphomimmetic states on the DBD. Figure 3A shows Phosphoserine has a formal charge of −2 at physiological pH, compared with −1 for glutamic acid, and thus engages in stronger electrostatic interactions with the cationic residues and sample two spatially distinct positions on that face, as observed in the distance analysis. Although both S295E and pS295 are localized near the DNA binding residues, stark difference in positional occupancy reflects a mechanistically important difference between S295E and the pS295 (Figure 3D), indicating a possibly allosteric mode of regulation. Interestingly, the basic residues on the DNA binding face that engage residue 295 in the phosphorylated states are the same residues proximal to DNA (Figures 3B-D). This suggests that the tethered NTR sits directly alongside the DNA binding surface and raises the possibility of physically hindering DNA accessibility.

EMSA data obtained using Ets2 proteins at different phospho-states confirm the above-mentioned possibility. Both S295E and phospho-Ets2 bind DNA with lower affinities (K_d_ ~0.9 µM for each protein, Figure 4A and B) compared to the unphosphorylated version of Ets2 (K_d_ ~0.4 µM). It is also observed that S295E binds DNA cooperatively whereas cooperativity in DNA-binding by unphosphorylated Ets2 or P-Ets2 were not apparent. with a plausible explanation of this unforeseen cooperativity lies in the partial allosteric perturbation of the DBD by S295E where its single charge tether perturbs the DNA-binding mode without abolishing it, requiring avidity contributions for occupancy. Comparable DNA-binding affinities of S295E and P-Ets2 strongly suggest that inhibition of DNA binding is primarily rendered through phosphorylation of S295, whereas additional phosphorylation might bestow other allosteric means to fine tune DNA binding properties like cooperativity. However, it is hard to confirm the exact nature of DNA binding cooperativity by pS295-Ets2 protein as phosphorylation of Ets2 cannot be restricted to a single site when treated with ATP and IKK2. Taken together, our biochemical and computational analyses provide a structural mechanism of phosphorylation mediated inhibition of DNA-binding by Ets2.

The regulatory paradigm presented here, where phosphorylation of the NTR induces intramolecular tethering to the DNA-binding face of the cognate structured domain, thus directly reducing DNA-binding affinity, expands the current paradigm of intrinsically disordered region-mediated regulation in ETS transcription factors [47–50]. In Ets2, the NTR-DBD interaction takes place directly on the DNA-binding face of the DBD, with the tethered NTR physically occluding DNA binding rather than allosterically redistributing electrostatics. This mechanistic divergence emphasizes the versatility of phospho-NTR regulation across ETS family members. More generally, the idea of disordered segment phosphorylation allosterically modulating structured binding domains via intramolecular contacts is in consistent with previous studies [47–50,52,53] and our work provides a structurally characterized example within the context of inflammatory kinase signaling through Ets2.

In sum, we elucidate a phosphorylation dependent regulation of DNA-binding activity of Ets2 by IKK2. Of note, similar phenomena are reported for Ets1 phosphorylation mediated by CaMKII. Taken together, we speculate that DNA-binding activities of ETS-transcription factors are dynamically controlled and fine-tuned by the cellular kinases in specific signaling contexts.

## Author contributions

Conceptualization: SP; Experimental design: SP, PK and DS; Investigation: DS, AH, PB, PRT (mass spectrometry), ARA (computational anayses); Data analysis: SP, DS, PK, SRC, PRT; Figure: DS, ARA, PK, SP; First draft: DS; Editing: SP, PK, DS.

## Acknowledgements

SP acknowledges funding from ANRF-IRHPA (IPA/2020/000414) and DBT-Wellcome Trust India Alliance Intermediate Fellowship to SP (IA/I/15/1/501852), and intramural core funding from Bose Institute; PK acknowledges funding support from ANRF(ANRF/ECRG/2024/000698/LS, ANRF/ARG/2025/007391/CS) and DBT (BioE3-mAb-BT/PR62183/MRNAR/169/14/2025), GoI. ARA thanks Ministry of Electronics and Information Technology (MeitY), Government of India for fellowship through IndiaAI scheme and AH thanks the University Grants Commission, Government of India for graduate research fellowship. Authors thank the proteomics facility at CSIR-CCMB for mass spectrometric data collection. Authors also thank Ms. Lahari Dasgupta for help with protein purification and Samrat Mitra for biochemical reagent and Prof. Siddhartha Roy for helpful discussion.

