## Supplementary material for "IKK2/β mediated phosphorylation of transcription factor Ets2 at site(s) distal to DNA binding domain negatively modulates its DNA binding activity": Materials and methods

***Materials***

All the chemicals used in this study were of molecular biology or analytical grade. Growth media included LB broth (#M1245, HiMedia) and LB agar (#M1151, HiMedia). The ‘HF’ versions of all restriction enzymes were purchased from New England Biolabs. Other enzymes and related reagents included Calf Intestinal Alkaline Phosphatase (CIAP) (#18009019, ThermoFisher Scientific), TaKaRa Taq™ DNA Polymerase (#R001C, Takara Bio), Q5® High-Fidelity DNA Polymerase (#M0491, NEB), Pfu DNA polymerase (#600382, Agilent Technologies), dNTP mixture (# 4030, Takara Bio), T4 DNA ligase (#EL0011, ThermoFisher Scientific), 10X T4 DNA ligase buffer (#B69, ThermoFisher Scientific). Antibiotics used were Ampicillin (MP-Biomedical, Cat no. 190148), Kanamycin (#K-120-25, GoldBio), and Chloramphenicol (#C-105-25, GoldBio).

Details of other reagents: Ni-NTA agarose (#30230, Qiagen), Agarose (#16500500, ThermoFisher Scientific), Adenosine-5’-triphosphate (ATP) (#A-081-25, GoldBio), Superdex 200 column (#28990944 or #90100137, Cytiva), 30% Acrylamide: bis-acrylamide 29:1 (cat. 1610156 Bio-Rad), Urea (#U-200-1,Gold Biotechnology), Tris (#648311-5KG, Merck), NaCl (#1.93606.5021,Merck), Glycerol (#1.07051.2521, Merck), 2-mercaptoethanol (#MB041-500ML, Himedia), EDTA (#T9191, Takara), SP HP column (#45-000-191,Cytiva), polydIdC (#P4929-10UN, Sigma Aldrich ), DTT (#D9779-25G, Sigma Aldrich), MgCl_2_ (CAS NO.[7791-18-6](https://www.merckmillipore.com/IN/en/search/7791-18-6?focus=products&page=1&perpage=30&sort=relevance&term=7791-18-6&type=cas_number)), HEPES (#194827, MP Biomedicals), Na3VO4 (#450243-50G, Sigma Aldrich), isopropyl-b-D-thiogalactopyranoside (#102101, MP Biomedicals), Imidazole (#56749-1KG, Sigma Aldrich ),phenylmethanesulfonylfluoride (#011687, Spectrochem), NH4HCO3 (09830-1KG, Sigma Aldrich), TCEP (#TCEP25,Gold Biotechnology), NaF (450022-25G, Sigma Aldrich), Tween20 (P9416-50ML, Sigma Aldrich), SDS (#1.94954.0521, Merck), β-Glycerophosphate (G9422), Iodoacetamide (IAA) (Catalog Number: VB1010, Promega ), rLysC (#V1671, Promega ), Trypsin Gold (#V5280,Promega), Calbiochem Inhibitor VII (CAS 873225-46-8)

***Methods***

**Cloning, expression and purification of proteins**

**Cloning:**

Ets2 FL(Human) construct was obtained in a mammalian backbone (pcDNA3) with N-terminal FLAG tag (Addgene cat #28121). At first Ets2 FL was cloned from pcDNA3 backbone to pFastBacHT B vector. Subcloning was done in bacterial expression vector pET28a from pFastBacHT B. Restriction enzymes used were BamHI and Xho I. Double digestion was done at 37℃ overnight. The vector was CIP treated and gel extracted. The Ets2 insert (1.4 kb) were also gel extracted from pFastBacHT B using the same set of enzymes. Ligation was set up in 1:3 ratio of the insert to the vector using T4 DNA Ligase at room temperature for 2hours. The ligation mixture was transformed into ultracompetent DH5α cells and the colonies were screened for the inserts using the same restriction enzymes and confirmed by sequencing.

Ets2 truncates was cloned in pColdI in BamHI and SalI sites. **Ets2 NTR-AI** spans 1-360amino acid (1080bp) and consists of the N-TERMINAL REGION and autoinhibition [1] domain (325-360amino acid) only. **Ets2 NTR** spans 1-325amino acid (975bp) and contains only the N-TERMINAL REGION. **Ets2 AI-DBD** stretches from 325-464 amino acid (417bp) and consists of both AI and ETS or DNA Binding (DBD) domain (360-464 amino acid). **Ets2 DBD** consists of the ETS or DBD (360-464 amino acid) only. Ets2 truncates were amplified by PCR using Q5 high-fidelity DNA polymerase (New England Biolabs) and subsequently subcloned into pColdI bacterial expression vector between the BamH I and Sal I restriction enzyme sites. The PCR products were digested with the mentioned enzymes and gel extracted. The backbone pCold I was also digested with the same enzymes followed by calf intestine alkaline phosphatase treatment and gel extracted. The Ets2 inserts and backbone were then set for ligation using T4 Ligase in 3:1 ratio at room temperature for 2hours. The ligation mixture was transformed into DH5α competent cells. The colonies were screened using NdeI and Sal I and finally confirmed by sequencing.

N-Terminal Ets2 truncates – **ΔN-84** and **ΔN-170** were cloned into pET28a bacterial expression vector. pET28a Ets2 FL was used as a template for PCR amplification using Q5 high-fidelity DNA polymerase (New England Biolabs) and subsequently digested using BamHI and XhoI enzymes. The vector was also digested with the same set of enzymes followed by CIP treatment. Both the insert and the vector were gel extracted. The Ets2 inserts and backbone were then set for ligation using T4 Ligase in 3:1 ratio at room temperature for 2hours. The ligation mixture was transformed into DH5α competent cells, and the colonies were screened for the inserts using the same restriction enzymes and confirmed by sequencing.

Site-directed mutagenesis was carried out for obtaining the S295E Ets2 phosphomimic mutant using pEt28a-Ets2 FL WT as the template. Overlap-extension PCR using complementary pair of primers containing desired base-mutation was used. Post-PCR amplification using *Pfu* Ultra DNA polymerase (Agilent Technologies), DpnI (New England Biolabs) was added to the reaction mixture to selectively digest the methylated parental DNA template. The digested products were transformed into *E. coli*, and plasmids from positive clones were Sanger-sequenced to confirm the presence of the intended nucleotide substitutions.

**Expression and purification:**

All the constructs were expressed in *E. coli* Rosetta2 (DE3) cells. NTR, NTR-AI, AI-DBD, ΔN-84, ΔN-170 and DBD were efficiently expressed in the soluble fraction. Cultures in LB were grown at 37 °C to OD_600_ ~ 0.4, induced with 0.3mM isopropyl-b-D-thio galactopyranoside (IPTG), and grown at 16 °C for ~ 18hours. Harvested cells were resuspended in buffer containing 50 mM Tris-HCl pH 8, 150 mM NaCl, 10 mM Imidazole, 5% glycerol, 5 mM 2-mercaptoethanol (BME), and 1 mM phenylmethanesulfonylfluoride (PMSF). Cells were lysed by sonication and centrifuged at 15000g for 1hour at 4 °C. After centrifugation the soluble supernatants were loaded onto pre-equilibrated Ni-NTA agarose beads for 1.5hours at 4 °C. The resin was washed with lysis buffer followed by second wash with wash buffer containing 20 mM Imidazole. Elution was done with 250 mM Imidazole containing buffer added in steps to create a gradient. Eluted fractions were analysed by SDS-PAGE. For NTR, NTR-AI, ΔN-84, ΔN-170 and AI-DBD, fractions containing purified protein were pooled and loaded onto a Superdex 200 gel filtration column (HiLoad 16/600 Superdex 200pg) in 50 mM Tris pH 8, 2% glycerol, 100 mM NaCl and 5 mM 2-mercaptoethanol (BME). Elution fractions containing DBD were dialysed in 20 mM K-phosphate pH-6, 50 mM NaCl, 5% glycerol,1 mM EDTA and 5 mM 2-mercaptoethanol (BME) overnight at 4 °C. The dialysed protein was then subjected to cation exchange chromatography using SP-HP column and eluted using high salt gradient. The elution fractions containing the protein was again dialysed in 20 mM K-phosphate pH-6.5, 150 mM NaCl, 5% glycerol,1 mM EDTA and 5 mM 2-mercaptoethanol (BME) and stored in -80 °C freezer.

Ets2 FL WT and S295E was expressed in *E. coli* Rosetta2 (DE3) cells. Cultures in LB were grown at 37 °C to OD_600_ ~ 0.4, induced with 0.3mM isopropyl-b-D-thiogalactopyranoside (IPTG), and grown at 16 °C for ~ 18hours. Harvested cells were resuspended in buffer containing 50 mM Tris-HCl pH 8, 150 mM NaCl, 10 mM Imidazole, 5% glycerol, 5 mM 2-mercaptoethanol (BME), and 1 mM phenylmethanesulfonylfluoride (PMSF). Cells were lysed by sonication and centrifuged at 15000g for 1hour at 4 °C. The supernatant was discarded and pellet washed with the same resuspension buffer containing additional 0.2% NP40. The pellet was then resuspended in 20 mM K-phosphate pH-8, 150 mM NaCl, 5% glycerol, 10 mM Imidazole, 5 mM 2-mercaptoethanol (BME), 1 mM phenylmethanesulfonylfluoride (PMSF) and 8 M Urea. The resuspended pellet in denaturing 8M urea buffer was centrifuged next day at 15000g for 1 hour at room temperature and the supernatant was used for purification. The supernatant was separated and was allowed to bind with Ni-NTA resin (equilibrated overnight in Resuspension buffer containing 8M) for 3-4hrs at room temperature. Wash 1 was same as Resuspension buffer and Wash 2 had 20mM Imidazole rest remaining same as Resuspension buffer. Elution was done using 250mM imidazole buffer (added in steps to create a gradient). The fractions containing the purified protein was set up for refolding in 20 mM K-phosphate pH-8, 150 mM NaCl, 5% glycerol and 5 mM 2-mercaptoethanol (BME). Size exclusion chromatography was performed with the refolded protein in 20 mM K-phosphate pH-8, 150 mM NaCl, 2% glycerol, 1 mM EDTA and 5 mM 2-mercaptoethanol (BME) and stored in -80 °C freezer.

IKK2 FL WT, IKK2 K44M and IKK2 AA was purified from Sf9 insect cells using respective recombinant baculoviruses following the protocol described in [2].

Protein concentration was determined by measuring absorbance at 280 nm using a Multiskan Sky High Microplate Spectrophotometer (Thermo Fisher Scientific) and calculated using the Beer-Lambert equation:

*A= ε x c x l*

where A is the absorbance at 280 nm, ε is the molar extinction coefficient of the protein, c is the protein concentration and l is the path length.

***In-vitro* radioactive kinase assay**

For every in-vitro radioactive kinase assay, a master mix was made as per the experimental set-up to minimize pipetting errors. Typically, 50-100 ng of purified kinase (catalytic amount) was used in each *in vitro* kinase reaction. 0.5-1μg of substrate (Ets2 FL WT, Ets2 truncates, S295E) was incubated with kinase for 30 mins or indicated time periods at 27˚C in presence of 20μM ATP with 0.1μCi of γ-P32-ATP in a reaction buffer containing 20mM HEPES pH 7.8, 100mM NaCl, 10mM MgCl_2_, 2 mM DTT, 10 mM Na_3_VO_4_, 10 mM NaF, 20 mM β-Glycerophosphate. Reaction was stopped by heating the samples at 100˚C for 5 mins after the addition of 1X-Laemmli buffer. Samples were resolved on SDS-PAGE. Gels were stained using Instant Blue Coomassie protein stain and the stained gels were dried using a gel drying apparatus (Bio-Rad Model 583) and exposed to a phosphor screen inside a cassette. Typically, the substrate band(s) were monitored for this purpose as kinase band(s) were not visible in catalytic amount. For radioactive kinase assays with K44M/IKK2 AA mutant, a higher amount of kinase was used (0.5-1μg), and the kinase band was visible (Fig. 1F). Extent of substrate phosphorylation was analyzed by phosphor-imaging (Typhoon, Cytiva). For the assays with inhibitors, IKK2 or IKK2: substrate was mixed with the respective inhibitors at concentrations shown in the respective figures for 30 minutes at room temperature in the kinase assay buffer prior to the addition of ATP or γ -P32-ATP. After the addition of ATP, reactions were allowed to proceed for specified time periods before analysing the results by autoradiography.[2-4]

**Electrophoretic mobility shift assay (EMSA)**

Fluorescent Cy5 labeled double-stranded 147bp DNA probes containing three binding sites for Ets2 were used in EMSA. EMSA buffer contained 50 mM Tris-HCl pH 7.5, 25 mM NaCl, 0.5 mM EDTA, 2 mM DTT, 5% glycerol, 0.1% Tween20, 5 mM MgCl_2_[1]. 25ng of poly dIdC was added to each reaction. Protein was incubated with Cy5-147bp DNA in EMSA buffer for 30mins at 25 ˚C. The reaction mixture was then loaded and resolved on a 6% native polyacrylamide gel in Tris Borate EDTA. The EMSA gels were visualised in Typhoon imager (Cytiva).

**Estimation of binding constant from EMSA**

For estimation of % of free DNA in each lane the following formulae was used:

% free DNA in lane x = (100 * free DNA intensity in lane x) / free DNA intensity in no protein sample

The approximate fraction of protein-bound DNA in each fraction can be obtained by subtracting the percentage of free DNA from 100.

% of bound DNA in lane x = 100 - % free DNA in lane x

For the graph fraction DNA bound and unbound were estimated:

Fraction unbound (DNA) = % free DNA in lane x / 100

Fraction bound (DNA) = 1 - Fraction unbound (DNA)

To determine the K_D_ of the DNA-protein interaction GraphPad Prism was used to plot the Fraction bound relative to the protein concentration in each sample. Data was fit with a non-linear regression model and Hill-Langmuir equation and a dissociation constant (K_D_) was obtained from this curve.[5]

**Mass Spectrometry**

To identify phosphorylation sites on Ets2 by IKK2, an in-vitro kinase assay was set up using 1mM non-radioactive ATP in kinase buffer containing 20mM HEPES pH 7.8, 100mM NaCl, 10mM MgCl_2_, 2 mM DTT, 10 mM Na_3_VO_4_, 10 mM NaF and 20 mM β-Glycerophosphate for 1hour at 27˚C. Reactions were then incubated with 0.1% SDS at 30 ˚C for 30 minutes. Buffer exchange was done in solution consisting of 50mM NH_4_HCO_3_, 8M urea and 2.5mM TCEP. The sample was then incubated at 37 ˚C for 45 minutes. After cooling to room temperature, Iodoacetamide (IAA) was added to a final concentration of 10mM and the reaction was incubated at room temperature for 1 hour in the dark to alkylate cysteine residues. Excess IAA was quenched by the addition of 5 mM DTT, followed by an additional 1-hour incubation at room temperature. Zeba Spin Desalting column was used to remove urea. To this rLysC was added at a 1:50 enzyme to substrate ratio (w/w) and incubated for 1 hour at 37 ˚C. Trypsin was reconstituted in 50 mM ammonium bicarbonate buffer and added to the samples at a 1:100 enzyme-to-substrate ratio (w/w). Trypsin was added in two steps: an initial addition was followed by incubation at 37°C for 2 hours with shaking, after which a second aliquot of trypsin (1:50 ratio) was added and the digestion was continued overnight at 37°C with shaking. The reactions were terminated by acidifying the samples with 0.1% formic acid at room temperature for 5 minutes to lower the pH below 3. The digested peptides were vacuum-dried using a SpeedVac (Thermo Scientific, Savant SPD111V), and phosphopeptides were selectively enriched using the High-Select Fe-NTA Phosphopeptide Enrichment Kit (ThermoFisher Scientific), following the manufacturer’s protocol. Peptides were analysed on an Orbitrap Exploris 240 mass spectrometer (Thermo Scientific) coupled to a nanoflow LC system (Easy-nLC II, Thermo Scientific). Samples were loaded onto a PepMap RSLC C18 nanocapillary reverse-phase column (75 μm x 25 cm, 2 μm particle size, 100 Å pore size) and separated using a 60-minute linear gradient of organic mobile phase consisting of 5% ACN with 0.1% formic acid (Buffer A) and 95% ACN with 0.1% formic acid (Buffer B). Raw MS data were processed using the MaxQuant computational proteomics platform (version 1.6.8) and searched against UniProt amino acid sequences corresponding to the protein of interest (UniProt ID: P15036). Phosphorylation of serine, threonine and tyrosine (STY) residues was specified as a variable modification in the search parameters.

**Sequence logo**

Protein sequences were taken from NCBI and aligned using Clustal omega and the IKK phosphorylation motif was visualized as a sequence logo using Weblogo 3 with default settings. [6,7]

**Structure modelling**

Structures of three different phosphorylation states namely S295 (wild type), S295E (phosphomimetic mutant), and pS295 (phosphorylated form) of ETS2 (UniProt[8] ID: P15036[9]) truncation, ΔN-80, were modeled to illustrate the role of the N-terminal region (NTR) in phosphorylation-dependent regulation. The S295 and S295E structures were modeled using AlphaFold3[10], whereas the pS295 structure was generated by replacing Serine at 295 in S295 by phosphoserine. The predicted models exhibited moderate overall confidence, with mean pLDDT[11] scores of 61.1 and 63.2 for the S295 and S295E models, respectively. However, the mean pLDDT scores for the structured DBD was found to be higher, 94.6 and 94.9 for S295 and S295E models respectively, with maximum value reaching to 98.5 for both models.

**System Setup for MD analyses**

The protein systems were built using AmberTools (v4.1)[12]. Standard amino acids were parameterized with the AMBER ff19SB force field[13], while phosphorylated residues were modeled with the phosaa19SB[14] extension that is compatible with ff19SB. Each system was solvated in a cubic TIP3P water box [15]with a minimum of 10 Å between the protein surface and the boundary of the box. 10, 11, and 12 Na^+^ counterions were added, respectively, to account for overall charge neutrality in S295, S295E, and pS295 systems. The generated AMBER topology and coordinate files were converted into GROMACS-compatible formats with ACPYPE[16].

**Molecular Dynamics Simulations**

MD simulations were performed using GROMACS 2024.5[17] on a Linux-based high-performance computing (HPC) cluster equipped with an AMD EPYC 9554P 64-core processor (128 hardware threads, 3.10 GHz). Energy minimization was performed using the steepest-descent algorithm for up to 500,000 steps or until the maximum force on any atom was below 1000 kJ mol⁻¹ nm⁻¹. Long-range electrostatic interactions were treated using the Particle Mesh Ewald (PME) method[18,19] with a real-space cutoff of 1.0 nm, while van der Waals interactions were truncated at 1.0 nm. Neighbor searching employed the Verlet cutoff scheme, and periodic boundary conditions were applied in all three dimensions.

Each system was equilibrated in two sequential phases namely constant volume and temperature (NVT) and constant pressure and temperature (NPT) equilibration. Initially, a 1 ns NVT equilibration was performed using the leap-frog integrator with a 2 fs integration timestep. The temperature was maintained at 300 K using the velocity-rescale (V-rescale) thermostat with a coupling constant (τ = 0.1 ps). Protein and non-protein atoms were coupled to separate temperature baths, and all Hydrogen-containing bonds were constrained using the LINCS algorithm[20]. Furthermore, a 1 ns NPT equilibration was performed using the same integrator and thermostat settings. Pressure was maintained at 1 bar using the Parrinello–Rahman barostat with isotropic pressure coupling (τ = 2.0 ps) and an isothermal compressibility of 4.5 × 10⁻⁵ bar⁻¹. Initial velocities for the NPT equilibration were inherited from the preceding NVT simulation.

For each system, the production run was for 500 ns in the NPT ensemble at 300 K and 1 bar using the same integration parameters. Long-range electrostatic interactions were calculated using the PME method with a real-space cutoff of 1.0 nm, a Fourier grid spacing of 0.16 nm, and fourth-order spline interpolation. Van der Waals interactions were truncated at 1.0 nm, and long-range dispersion corrections were applied to both energy and pressure. Trajectory coordinates were saved every 10 ps for subsequent analyses.

**Trajectory Analysis**

Trajectories were corrected for periodic boundary conditions and aligned to the DBD by least squares fitting to remove overall translational and rotational motion before analysis. The resulting trajectories were analyzed for structural stability, residue-level flexibility, conformational sampling, spatial occupancy of residue 295, electrostatic surface properties of the DBD, and phosphorylation-dependent intramolecular interactions between residue 295 and basic residues of DBD. We used GROMACS[17], MDAnalysis[21], ChimeraX[22,23], Matplotlib[24], and *in-house* Python[25] scripts for analyses.

The conformational sampling of residue 295 was characterized using three-dimensional residue occupancy maps generated from the production trajectories. These maps were used to quantify the spatial distribution and preferred conformational states sampled by residue 295 relative to the DBD.

Electrostatic interactions between residue 295 and the DBD were examined by calculating the minimum distances between residue 295 and positively charged lysine and arginine residues of the DBD. Distance distributions and time-resolved interaction maps were generated to identify persistent intermolecular contacts throughout the simulations.

Electrostatic surface potentials of the ETS2 DBD were calculated using the Adaptive Poisson–Boltzmann Solver (APBS)[26,27] and visualized in ChimeraX. Electrostatic surfaces were compared with residue-occupancy distributions to assess the spatial relationship between residue 295 and the DBD's charged regions.
